# Altered mitochondrial respiration is associated with loss of nuclear-encoded OXPHOS genes in parasitic broomrapes

**DOI:** 10.1101/2025.02.10.637464

**Authors:** Liming Cai, Robert K Jansen, Justin C Havird

## Abstract

Parasitic plants, characterized by their dependency on host organisms for nutrients, have displayed far-reaching alterations in physiology and genetics. While significant gene losses and relaxed selection have been documented in the nuclear and plastid genomes, how parasitism impacts the molecular evolution and function of mitochondria has remained controversial. One of the main culprits hindering our understanding in this area is the lack of knowledge on nuclear-encoded mitochondrial-targeted genes (N-mt), which encode most mitochondrial oxidative phosphorylation (OXPHOS) proteins. By conducting a comprehensive survey of N-mt genes across angiosperms, we demonstrated significant gene losses and horizontal transfers associated with relaxed selection unique to holoparasitic Orobanchaceae. These putative losses and transfers have the potential to affect mitochondrial function directly and cause mitonuclear incompatibility because of breakdown between co-evolved protein complexes from mitochondrial and nuclear genomes. Our physiological assessments using high-resolution respirometry revealed that despite genetic alterations, holoparasitic Orobanchaceae maintained robust OXPHOS function but relied more on the fully nuclear-encoded succinate dehydrogenase (complex II). Our results document the first example of significant and biased gene loss in nuclear-encoded mitochondrial OXPHOS genes in parasitic plants, expanding on previous studies focused on mitochondrial-encoded genes and elucidating the mechanisms underlying the preservation of OXPHOS function despite genomic reduction.

## INTRODUCTION

Mitochondria fuel energy metabolism for nearly all eukaryotes including plants. Chemical energy is converted to ATP via a series of orchestrated oxidative phosphorylation enzymes (OXPHOS) and accessory proteins that transport electrons, pump protons, and ultimately convert ADP to ATP. In most eukaryotes, this process is carried out by five core protein complexes I–V (CI-CV) on the inner mitochondrial membrane, including NADH dehydrogenase (CI), succinate dehydrogenase (CII), cytochrome c reductase (CIII), cytochrome c oxidase (CIV), and ATP synthase (CV). These enzymes vary greatly in size and genetic makeup, with many containing chimeric assemblies of mitochondrial and nuclear-encoded proteins. Consequently, these chimeric OXPHOS complexes are not only hotspots of cytonuclear coevolution, but also vulnerable targets of genetic incompatibility (Sloan *et al*., 2017, 2018; Havird *et al*., 2019*a*). Such incompatibilities come with strong fitness costs and can play a major role in creating reproductive barriers in yeasts, fishes, and plants (Lee *et al*., 2008; Luo *et al*., 2013; Moran *et al*., 2024). A potentially more widespread and less well-documented consequence of cytonuclear conflict is compromised or altered mitochondrial respiration (Havird *et al*., 2019*b*; Weaver *et al*., 2020). In plants, the electron transfer chain is further branched with alternative electron entry and exit via the alternative NAD(P)H dehydrogenases (NDs) and alternative oxidase (AOX). These complexes can alter OXPHOS function in response to environmental stressors and even compensate for OXPHOS dysfunction caused by cytonuclear conflict (Siedow and Girvin, 1980; Vanlerberghe, 2013; Sweetman *et al*., 2019). For example, the extremely high mitochondrial substitution rates of *Silene* (Caryophyllaceae) have led to lower efficiencies in chimeric OXPHOS enzymes such as CI and CIV, along with increased reliance on alternative NDs and AOX for mitochondrial respiration (Weaver *et al*., 2020).

Cytonuclear conflict may stem from the proliferation of incompatible mitochondrial mutations that can be attributed to changes in population dynamics, DNA repair machinery, mutational processes, life history strategies, or a combination of these phenomena (Havird *et al*., 2017; Zwonitzer *et al*., 2024). The evolution of parasitism, in particular, may trigger abrupt changes in energy metabolism that lead to shifts in how chimeric versus alternative OXPHOS enzymes are used. The most well-characterized and enthralling examples come from parasitic microbial eukaryotes — A wide range of mitochondrial functions have been documented from fully aerobic organelles to mitosomes stripped of OXPHOS and restricted to Fe-S biosynthesis (Hjort *et al*., 2010; Santos *et al*., 2018; John *et al*., 2019; Mathur *et al*., 2021). In at least one case the obligate symbiont *Monocercomonoides* has lost its mitochondria completely (Karnkowska *et al*., 2016). In photosynthetic organisms, comparable mitochondrial alterations have been documented especially among parasitic algae (Reyes-Prieto *et al*., 2002; Hancock *et al*., 2010; Smith *et al*., 2010; Smith and Lee, 2014). Among flowering plants, the most well-studied example comes from the European mistletoe *Viscum* (Santalaceae). Their CI genes have been lost completely along with highly diverged sequences for the other mitochondrial genes, which is indicative of relaxed selection on mitochondrial DNA (Petersen *et al*., 2015; Skippington *et al*., 2015). These changes have diminished mitochondrial respiration function fivefold, leading to an increased dependence on alternative pathways via NDs and AOX (Maclean *et al*., 2018).

The exciting discoveries in mistletoes have sparked a rush of investigations on parasitic and mycoheterotrophic plants. Yet none showed comparable levels of gene loss (Zervas *et al*., 2019). Instead, more subtle genomic changes have been found in mostly non-coding regions, including structural rearrangements, horizontal gene transfers, repeat expansion, nucleotide composition bias, and substitution rate elevation (Xi *et al*., 2013; Fan *et al*., 2016; Sanchez-Puerta *et al*., 2017; Petersen *et al*., 2019; Shtratnikova *et al*., 2020; Yu *et al*., 2023). These genetic alterations showed limited impacts on mitochondrial respiration, even in Balanophoraceae, where more than 80% of the mitochondrial-encoded protein-coding genes are replaced by horizontally transferred genes (Gatica-Soria *et al*., 2024). Thus, many have concluded that the association between parasitism and functional genomic degradation is restricted to the plastid and nuclear genomes and not mitochondria (Fan *et al*., 2016), with mistletoes being an exception rather than the rule.

However, previous studies neglected the majority (77%) of OXPHOS proteins encoded by nuclear genes along with an even larger repertoire of mitochondrial assembly factors and genes involved in DNA replication, repair, and recombination (RRR) that may impact the overall integrity of the mitochondrial genome and function (Sloaan, 2015; Smith and Keeling, 2015). Furthermore, functional verifications on OXPHOS respiration are outstandingly lacking except for *Viscum* and very recently, in Balanophoraceae (McLean *et al*., 2018; Gatica-Soria *et al*., 2024). The impact of parasitism on mitochondrial respiration cannot be fully understood without a comprehensive survey of the content of OXPHOS genes in both the mitochondrion and nucleus, ideally combined with thorough functional assessments.

To remedy this knowledge gap, we used comparative genomics and high-resolution respirometry to study the link between nuclear-encoded mitochondrial-targeted gene (N-mt) content, mitochondrial respiration, and parasitism in the flowering plant family Orobanchaceae (the broomrapes). These ca. 2,000 species span the entire spectrum of plant parasitism from free-living species to holoparasites that rely completely on hosts for water and nutrients. In particular, the transition to holoparasitism has evolved three times independently (McNeal *et al*., 2013), providing a unique comparative framework to investigate the link between life history strategy and organellar function. In contrast to the well-characterized plastid genome degradation of Orobanchaceae (e.g., (Wicke *et al*., 2016)), mitochondria have received less attention. Mitochondrial genomes from approximately a dozen Orobanchaceae hemiparasites and holoaparasites have a complete and canonical set of core genes with only slightly elevated substitution rates in some lineages (Fan *et al*., 2016; Zervas *et al*., 2019). Widespread horizontal gene transfer has been reported in mitochondria, but cases are restricted to non-coding regions such as the mobile group I intron of the *cox1* gene (Fan *et al.,* 2016). Yet given their fundamental changes in energy metabolism, the overall trend of genomic degradation in parasites, together with their strongly host-dependent reproductive strategy that constantly bottlenecks the population (Cai *et al*., 2021; Cai, 2023), we hypothesize relaxed selection in the mitochondrial DNA of parasitic Orobanchaceae. This may lead to changes as extreme as the entire loss of CI in *Viscum*, or more subtle signatures such as accelerated rates in N-mt genes.

Here, we conducted a comprehensive survey of N-mt genes across angiosperms, focusing on species in Orobanchaceae spanning parasitic life history strategies. We demonstrated frequent losses and significant relaxed selection of N-mt genes unique to one holoparasitic tribe (Orobancheae sensu Fischer (Fischer, 2004)) that are otherwise rare in flowering plants. These included both OXPHOS and mitochondrial RRR genes. To explore the functional impact of these gene losses, we applied high-resolution respirometry to isolated mitochondria from representative species to generate fine-scale mitochondrial respiration data at the protein level. We found that parasitic lineages with extensive N-mt gene loss tend to have increased reliance on nuclear-encoded CII, alternative NDs, and AOX, although CI and other chimeric OXPHOS complexes maintained normal function despite gene losses.

## MATERIALS AND METHODS

### Comparative survey of N-mt genes

Homologs of N-mt genes were identified in two steps. First, a broad survey of N-mt genes across angiosperms was conducted using an orthogroup database from our previous studies (Cai *et al*., 2021). This database contains 23,151 orthogroups from 28 angiosperm species inferred by the similarity-based orthology clustering program OrthoFinder v2.2.7 (Emms and Kelly, 2015) (electronic supplementary material, table S1). These species have high quality reference genomes spanning from early-diverging *Amborella* to representative species in rosids and asterids. To retrieve N-mt genes, N-mt homologs of *Arabidopsis thaliana* reported by Meyer *et al*. (Meyer *et al*., 2019) and Zhang *et al*. (Zhang *et al*., 2016) were used as baits to identify the corresponding orthogroups. We included 61 nuclear-encoded OXPHOS proteins, 7 mitochondrial ribosomal proteins, and 26 organellar RRR proteins (18 dual-targeted to mitochondria and plastids, plus 8 specific to mitochondria). To mitigate false negatives stemming from BLAST algorithms or annotation errors, we further verified all absences by two approaches: (1) We used an alternative similarity-based search tool — the profile hidden Markov implemented in HMMER v3.3.2 (Mistry *et al*., 2013) — to identify any potential homologs in coding sequences with an e-value threshold of 1e-30; (2) We then searched the entire genome sequences from these 26 reference taxa using BLASTN to confirm absence or pseudogenization. Whenever possible, multiple versions of genome assemblies and annotations from the same species or close relatives from the same genus were consulted (electronic supplementary material, table S1).

Second, a focused search of N-mt genes in Orobanchaceae was conducted by compiling a database using published genomes and transcriptomes from 15 species (electronic supplementary material, table S1). These included one free-living species (*Lindenbergia philippensis)*, seven hemiparasites, and seven species that represent two independent origins of holoparasitism (figure 1). We then applied BLASTN and HMMER to identify candidate N-mt genes as described previously. Due to the incomplete nature of transcriptomes, gene loss was determined conservatively, with any orthologous fragments counted as present (except when frameshifts or premature stop codons were confirmed manually). When possible, sequence data from multiple individuals of the same species were used to verify putative gene losses in *Epifagus virginiana* (two RNA samples), *Aphyllon fasciculatum* (two RNA samples), and *Lindenbergia philippensis* (one genome and two RNA samples; electronic supplementary material, table S1).

**Figure 1.**
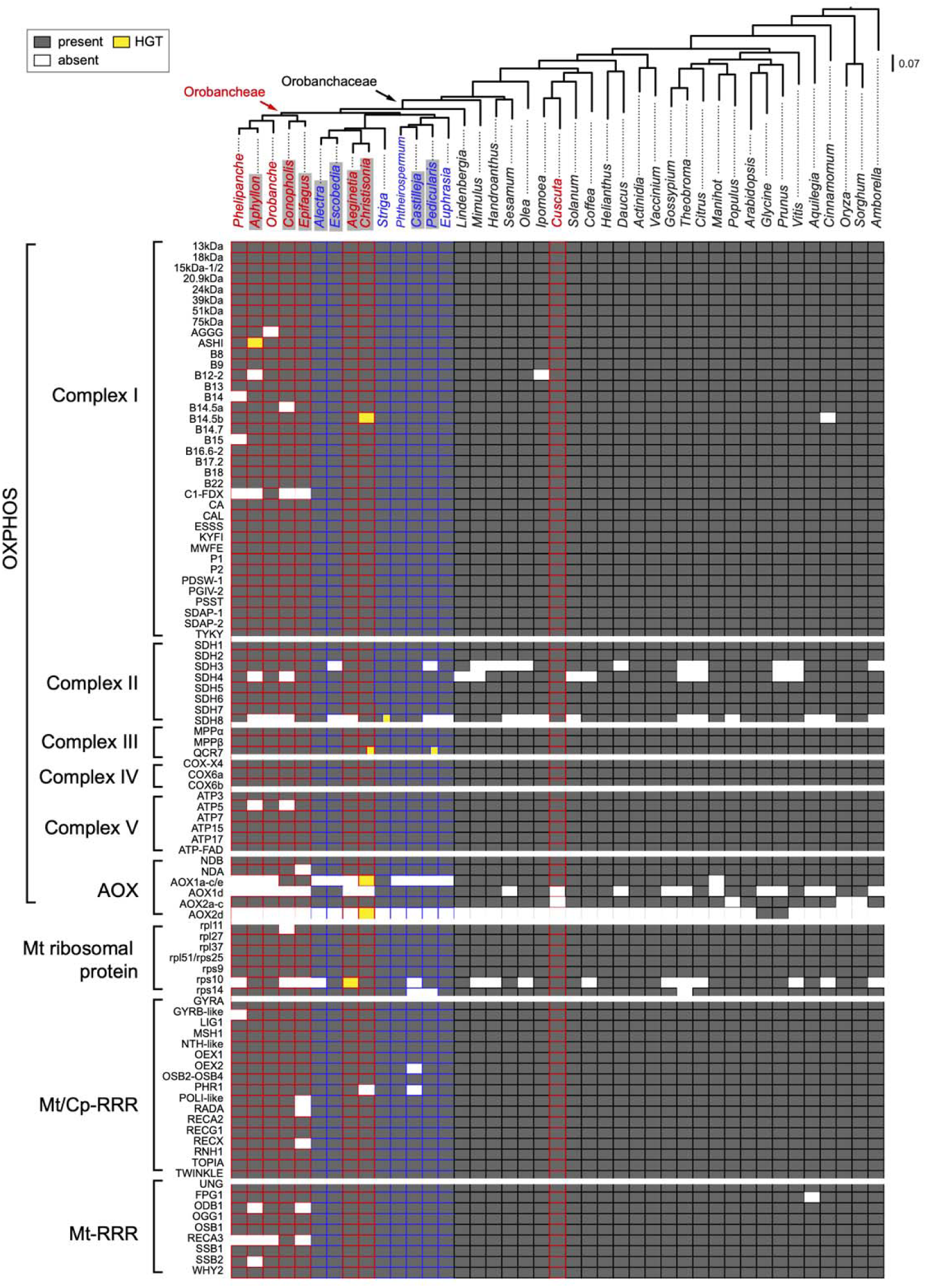
Presence and absence matrix of selected nuclear-encoded mitochondrial (N-mt) enzymes shows major losses in the holoparasitic tribe Orobancheae. The maximum likelihood phylogenetic tree of representative angiosperms is inferred from a concatenated DNA alignment of all N-mt genes in IQ-TREE. Holoparasitic lineages are highlighted in red and hemiparasitic lineages are highlighted in blue. Species with transcriptome data are marked with gray shading. Dark grey cells indicate complete or partial presence of the sequence; white cells indicate absence; yellow cells indicate putative horizontal gene transfer; hybrid cells indicate coexistence of both native and horizontally transferred sequences.

Finally, candidate orthologous sequences from Orobanchaceae were combined with the angiosperm dataset. A phylogeny-based approach was implemented to remove non-orthologous sequences. To accomplish this, individual gene trees were inferred with DNA alignments, and the maximum inclusion algorithm described in Yang and Smith (Yang and Smith, 2014) was applied. This algorithm iteratively cuts out the subtree with the highest number of taxa without taxon duplication. It also outputs a one-to-one ortholog dataset that can be directly used for species tree inference. Details on DNA alignment and phylogeny inference are described in the section ‘Species tree reconstruction’ below. All scripts used in this study are available on GitHub (https://github.com/lmcai/Orobanchaceae_O2K).

### Species tree reconstruction

To infer a species tree for comparative analyses, we aligned the protein sequences of one-to-one orthologs using the E-INS-i algorithm implemented in MAFFT v7.453 (Katoh and Standley, 2013). These protein alignments were then back translated to codons with pal2nal v14 (Suyama *et al*., 2006). Maximum Likelihood phylogenies were inferred for both individual genes and the concatenated codon alignments using IQ-TREE v2.2.2.6 (Minh *et al*., 2020). A gene-by-gene partition was used for the concatenation analysis and the best substitution model was determined by the built-in ModelFinder program (Kalyaanamoorthy *et al*., 2017). Branch support was assessed by performing 1000 ultrafast bootstrap replicates (UFBP).

### Verification of missing N-mt genes

In addition to applying alternative search algorithms and multiple assembly or annotation versions, we further examined whether data quality or sequence divergence may lead to observed patterns of missing genes. First, genome and transcriptome quality were assessed using BUSCO v5.7.0 based on the seed plant (embryophyta_odb10) and the eukaryote (eukaryota_odb10) databases (Simão *et al*., 2015). Both databases were applied because the seed plant BUSCO database is enriched in photosynthesis-related genes that are prone to loss in parasitic plants (Cai *et al*., 2021). Therefore, the eukaryote database is more accurate than the seed plant database for quality control in this case. Second, we examined the pairwise sequence divergence between *Arabidopsis* and other angiosperms to test whether missing genes may arise artificially from extreme sequence divergence. We applied the dist.dna function from the R package ape (Paradis and Schliep, 2019) to calculate both the proportion of nucleotide difference (model = “raw”) as well as the sequence divergence under the F84 substitution model (model = “F84”). The concatenated DNA alignment of N-mt genes for species tree inference was used for this calculation.

Finally, we used phylogenetic ANOVA to evaluate whether parasitic or holoparasitic Orobanchaceae demonstrated significantly higher proportions of missing BUSCOs or higher sequence divergence compared to free-living angiosperms. The species tree inferred above and the phylANOVA function in the R package phytools (Revell, 2012) were used. To further evaluate the significance of N-mt gene loss in Orobanchaceae while accounting for data quality, we conducted a Fisher’s exact test with *p*-values corrected by the ratio of missing eukaryote BUSCOs.

### Molecular evolution

To test for relaxed selection on N-mt genes in parasites, we grouped genes into nine functional classes (figure 1) and then applied the RELAX method to test for relaxed selection in HyPhy v2.5.33 (Wertheim *et al*., 2015; Kosakovsky Pond *et al*., 2020). RELAX compares the ω value (d_N_/d_S_) in the foreground branches to the background branches in a reference species tree. If the foreground branches are under relaxed selection, their ω values are expected to converge towards the neutral value 1 across all rate categories and vice versa for intensified selection (Wertheim *et al*., 2015). To apply this method, we subsampled the codon alignment to include a single sequence per species to facilitate the function-based concatenation. Potential horizontal gene transfers nested outside Orobanchaceae were removed prior to downstream analyses (see Results below). When multiple copies were represented for a species, we chose the one showing minimal root-to-tip distance to facilitate concatenation. The species tree inferred above was used as the reference. We then ran RELAX under 3 rate categories, the GTR substitution model, and constant synonymous substitution rates across sites. We tested two evolution models where either the holoparasitic tribe Orobancheae alone or Orobancheae plus the other holoparasitic lineage *Aeginetia*+*Christisonia* were selected as foreground test branches (figure 1). The goodness of fit of these two models was evaluated using the likelihood ratio test.

### Taxon sampling for respirometry assessment

To explore the functional impact of OXPHOS gene loss, we assessed mitochondrial respirometry in representative species of Orobanchaceae. This experiment requires fresh tissue and access to a molecular lab immediately after collection. Moreover, the scarce distribution and ephemeral above-ground phenology of many Orobanchaceae further restricted our sampling options. We therefore selected fifty individuals from nine species adjacent to our laboratory facilities in Austin, TX and Cambridge, MA during two field seasons from 2022 to 2023 (electronic supplementary material, table S2). All four holoparasites — *Aphyllon uniflorum*, *Orobanche minor*, *Phelipanche ramosa*, and *Conopholis americana* — are members of the tribe Orobancheae, which have experienced extensive OXPHOS gene loss (see Results). The other five hemiparasitic and free-living species served as references where OXPHOS genes are under stronger purifying selection. These plants were carefully excavated from the soil, sometimes with their hosts to avoid stress. To account for variation in tissue types and developmental stages, we exclusively sampled floral tissues when possible. Due to the variation in phenology and the large quantity of required tissue (1 g per experiment), leaves were sampled for *Aureolaria pedicularia* (electronic supplementary material, table S2).

Besides the nine species presented in this study, respirometry assays were also performed on six hemiparasitic and free-living species including *Belladia trixago, Melampyrum lineare*, *Aureolaria flava*, *Triphysaria versicolor, Rehmannia glutinosa*, and *Rehmannia grandiflora*. However, experiments were inconclusive for these taxa, given that respiration rates did not significantly increase when the substrate ADP was added (see below). We attribute these negative results to organism- and tissue-specific metabolic profiles that might inhibit the standard respirometry assay, rather than a real loss of mitochondrial function and do not present respirometry data for these species here.

### Mitochondrial isolation and respirometry

Wild plants were harvested and preserved on ice before being transferred to the lab. Respirometry assays followed a well-established protocol in our lab that has been applied to a diverse array of plant and animal systems including *Silene,* crustaceans, nematodes, and mayflies (Havird *et al*., 2019*b*; Weaver *et al*., 2020). A detailed protocol is provided in electronic supplementary material, note S1. Briefly, 1 g of floral or leaf tissue was minced and ground in an ice-cold mitochondrial isolation buffer following Havird *et al*. (Havird *et al*., 2019*c*). Intact mitochondria were then isolated using differential centrifugation and protein content of mitochondrial isolates was quantified. Subsequently, we used the Oroboros O2K high-resolution respirometry system (Innsbruck, Austria) with a substrate-uncoupler inhibitor-titration protocol to quantify seven different mitochondrial respiration states in each sample (electronic supplementary material, figure S1). We recorded the maximum respiration rate observed prior to the addition of any inhibitors, the maximum rate observed during the experiment, as well as several internally normalized statistics (see below). The calculated ADP-driven respiration rate (in nmol O_2_ s^-1^ mL^-1^ g^-1^) was achieved after the addition of succinate. The maximum respiration rate was normalized by the protein content and we thus excluded three species (*Aphyllon uniflorum*, *Conopholis americana,* and *Aureolaria pedicularia*) from maximum respiration measurements because they lacked protein quantification due to the inaccessibility of lab equipment in the field (electronic supplementary material, table S2).

### Statistical analyses

To examine the relative contributions of OXPHOS enzymes, six flux control factors (FCFs) were calculated (electronic supplementary material, table S3). Flux control factors are internally normalized by the ratio of respiration rate before and after the addition of electron donors or inhibitors to specific OXPHOS components (Pesta and Gnaiger, 2012). Two of the FCFs are related to CI function: OXPHOS efficiency (analogous to the respiratory control ratio (Jacoby *et al*., 2015)), and a CI FCF induced by adding the CI inhibitor rotenone. We also calculated FCFs that described CII, CIV, and the nuclear-encoded external NAD(P)H dehydrogenases (DH_ex_) and AOX. Linear mixed-effects (LME) models were implemented using the lme function from the R package nlme to test for statistical significance in differences in respiratory flux control (version) *et al*., 2024). Species were classified into two groups: with and without N-mt gene loss (i.e., free-living+hemiparasitic vs. holoparasitic). To control for multiple observations within a species, we included species as a fixed factor and accession as a random factor. We log-transformed the CIV FCF to meet the assumption of normality. We also tested for correlations among FCFs and maximum respiration rate by fitting LME models.

## RESULTS

### Conserved N-mt gene content across angiosperms

The availability of high-quality genomes in reference taxa across the angiosperm tree of life has greatly facilitated the refinement of comparative gene surveys. All reference angiosperm species outside Orobanchaceae exhibited a conventional and nearly complete set of N-mt OXPHOS genes. Exceptions existed in CII, where nuclear-encoded *SDH3* and *SDH4* were absent or recovered as relic, highly reduced gene fragments in 10 (38%) and 7 (27%) species, respectively (figure1). These absences are expected because both genes are still primarily encoded by mitochondria in many angiosperms with frequent mito-to-nuclear gene transfers in others (Palmer *et al*., 2000; Huang *et al*., 2019; Mower, 2020). Therefore they are not detectable in reference genomes that only include the nuclear sequences. The accessory subunit *SDH8* of CII was also frequently absent, but this is likely to be a result of its small size (46 amino acids in *Arabidopsis*) making it challenging to identify bioinformatically (Huang *et al*., 2019). Among the other four major complexes, CI is the largest, consisting of 52 proteins, but we found only two spontaneous losses of accessory proteins *B14.5b* in *Cinnamomum* (Lauraceae) and *B12* in *Ipomoea* (Convolvulaceae). For the AOX complex, one loss of AOX1 was found in *Manihot* (Euphorbiaceae) and three losses of the more dispensable and stress-related AOX2 were found in monocots, *Populus* (Salicaceae), and *Cuscuta* (Convolvulaceae). Beyond OXPHOS, the ribosomal and mitochondrial RRR proteins were universally present except for the *rps10* ribosomal protein, where putative losses were reported in 10 species (figure 1).

Despite the consistent presence of these N-mt genes, angiosperm lineages possessed variable numbers of homologs due to historical and ongoing duplication, loss, and transfer. Such copy number variations were lineage- and gene-specific. For example, the average gene copy number was three times higher in the blueberry *Vaccinium corymbosum* (n = 3.9) compared to all other species (n=1.3). This is likely a result of the combined effects of multiple rounds of whole genome duplications as well as tandem gene duplications in *Vaccinium* (Larson *et al*., 2020; Wang *et al*., 2020). Similarly, species in asterids and rosids had multiple copies of *MPPα,* coinciding with the core eudicot hexaploidy event (electronic supplementary material, data S1; (Jaillon *et al*., 2007)).

### Loss of N-mt genes is associated with relaxed selection in Orobanchaceae

N-mt gene losses in Orobanchaceae exhibited a strong lineage-specific pattern that was restricted to the holoparasitic tribe Orobancheae (figure 1). This pattern was not characteristic of all holoparasitic lineages, because we found limited gene loss in the lineage consisting of *Aeginetia* and *Christisonia*. We made substantial efforts to verify this lineage-specific gene loss pattern — First, data quality is unlikely to explain this pattern because Orobanchaceae did not show more missing eukaryote BUSCOs compared to other angiosperms (phylANOVA *p-*value = 0.266; electronic supplementary material, table S1). However, there were more fragmented eukaryote BUSCOs in Orobanchaceae (phylANOVA *p-*value = 0.003) due to the nature of transcriptome. There were also more missing seed plant BUSCOs in Orobanchaceae due to the evolution of parasitism (phylANOVA *p-*value = 0.001; (Cai *et al*., 2021)). Yet our conservative approach of counting fragmented N-mt genes should accommodate this data quality issue. Furthermore, the holoparasitic tribe Orobancheae demonstrated significantly more losses in N-mt genes when corrected for the background missing genes using 10000 empirical permutations in Fisher’s exact test (adjusted *p-*value < 1e-4). Second, Orobanchaceae did not show higher sequence divergence with *Arabidopsis* compared to other angiosperms (electronic supplementary material, table S4). The average pairwise sequence divergence between Orobanchaceae and *Arabidopsis* is 0.32 for raw nucleotide difference and 0.42 under the F84 substitution model, both of which are comparable to other angiosperms (phylANOVA *p-*value 0.52 and 0.55, respectively). Finally, the strong phylogenetic signal of missing N-mt genes, confined to the tribe Orobancheae, strongly supports this finding. In contrast, a full set of N-mt genes were readily identifiable in other hemi- and holoparasitic species with comparable data quality to that in Orobancheae. Losses within Orobancheae were also validated by two high-quality genomes from *Phelipanche aegyptiaca* and *Orobanche cumana.* In two species *Epifagus virginiana* and *Aphyllon fasciculatum*, N-mt losses were supported by multiple RNA samples from different individuals (electronic supplementary material, table S1).

Within tribe Orobancheae, gene losses were most significant in CI and CV, but were not remarkable in CII, CIII, and CIV. Most of the losses in CI were restricted to accessory proteins and have taken place several times independently in species such as *Phelipanche aegyptiaca*, *Orobanche cumana*, *Conopholis americana*, or their common ancestors (figure 1). These included membrane proteins *B12-2* and *AGGG* as well as peripheral proteins *B14.5a* and *C1-FDX* (Subrahmanian *et al*., 2016). Moreover, we also found two rare losses in CV located in *ATP5* which encode the OSCP subunit of the peripheral flexible hinge of the ATP synthase (Zancani *et al*., 2020). Beyond OXPHOS enzymes, the holoparasitic Orobancheae also only had 80.7% of the organellar and mitochondrial-specific RRR genes compared to other angiosperms.

Although a nearly complete set of N-mt genes was present in *Aeginetia* and *Christisonia*, this lineage is a hotspot for horizontal gene transfer (figure 1; electronic supplementary material, data S1). In at least eight cases, host-derived N-mt genes either replaced or were found alongside native copies. These include three repeated transfers of OXPHOS genes from Poaceae to *Christisonia* or *Aeginetia* (81–98 UFBP in IQ-TREE); one from Poaceae to *Pedicularis* (82 UFBP); one from Apiaceae to *Striga* (78 UFBP); one from Rosaceae to *Christisonia* (99 UFBP); and one from Asteraceae to *Aphyllon* (100 UFBP; electronic supplementary material, data S1). All putative gene transfers were restricted to OXPHOS genes and none were identified among the RRR genes. No transfer was found in *Cuscuta australis* (Convolvulaceae), which is a holoparasitic plant from the morning glory family.

Both gene losses and horizontal transfers are signatures of relaxed selection as supported by the results from RELAX analyses. Tribe Orobancheae has experienced significantly relaxed selection in CI, CII, CIII, and CV compared to all other angiosperms (*p-*value <0.05; electronic supplementary material, table S5). Significantly intensified selection was detected in the RRR genes that remained in Orobancheae (*p-*value = 0.0061), while the CIV, AOX, and ribosomal genes do not show significant differences in selective pressure in Orobancheae. An alternative model where relaxed selection took place in both tribe Orobancheae and the holoparasitic *Aeginetia* +*Christisonia* clade was not favored based on the likelihood ratio test (ΔLog(L) = 0.30, Chi-square *p-*value >0.05).

### Evolution of AOX in Orobanchaceae

Given its potential role in mitochondrial functional rescue, we investigated the evolution of AOX genes in Orobanchaceae, which are responsible for alternative respiratory pathways. We identified four clades: AOX1a–c/1e, AOX1d, AOX2a–c, and AOX2d in the two subfamilies AOX1 and AOX2, as suggested by previous investigations (figure 2; electronic supplementary material, data S1; (Costa *et al*., 2014)). In free-living angiosperms, the AOX1a–c/1e and AOX2a-c clades are nearly universally present except that AOX2a-c is absent in monocots. The AOX1d and AOX2d clades, on the other hand, are more dispensable and are hypothesized to originate from convergent evolution for stress response in plants (Costa *et al*., 2014). In Orobanchaceae, widespread losses are identified in the highly conserved AOX1a-c/1e clade but the AOX1d is relatively conserved (electronic supplementary material, data S1). AOX1a-c/1e is only recognizable in *Conopholis* +*Epifagus*, *Striga*, *Lindenbergia,* as well as *Christisonia,* the last of which is a result of gene transfer from likely Convolvulaceae (86 UFBP). This is in stark contrast with other angiosperm lineages where AOX1a-c/1e was identified in all species except *Manihot.* On the other hand, AOX2a-c is present in all Orobanchaceae with a few losses outside this family in *Cuscuta* (Convolvulaceae), *Populus* (Salicaceae), and monocots. The distinct AOX2d clade, though, was restricted to a few rosid families such as Fabaceae and Rosaceae. It is not present in any Orobanchaceae species except in the gene transfer from a Rosaceae clade including *Prunus* and *Fragaria* to *Christisonia* (electronic supplementary material, data S1).

**Figure 2.**
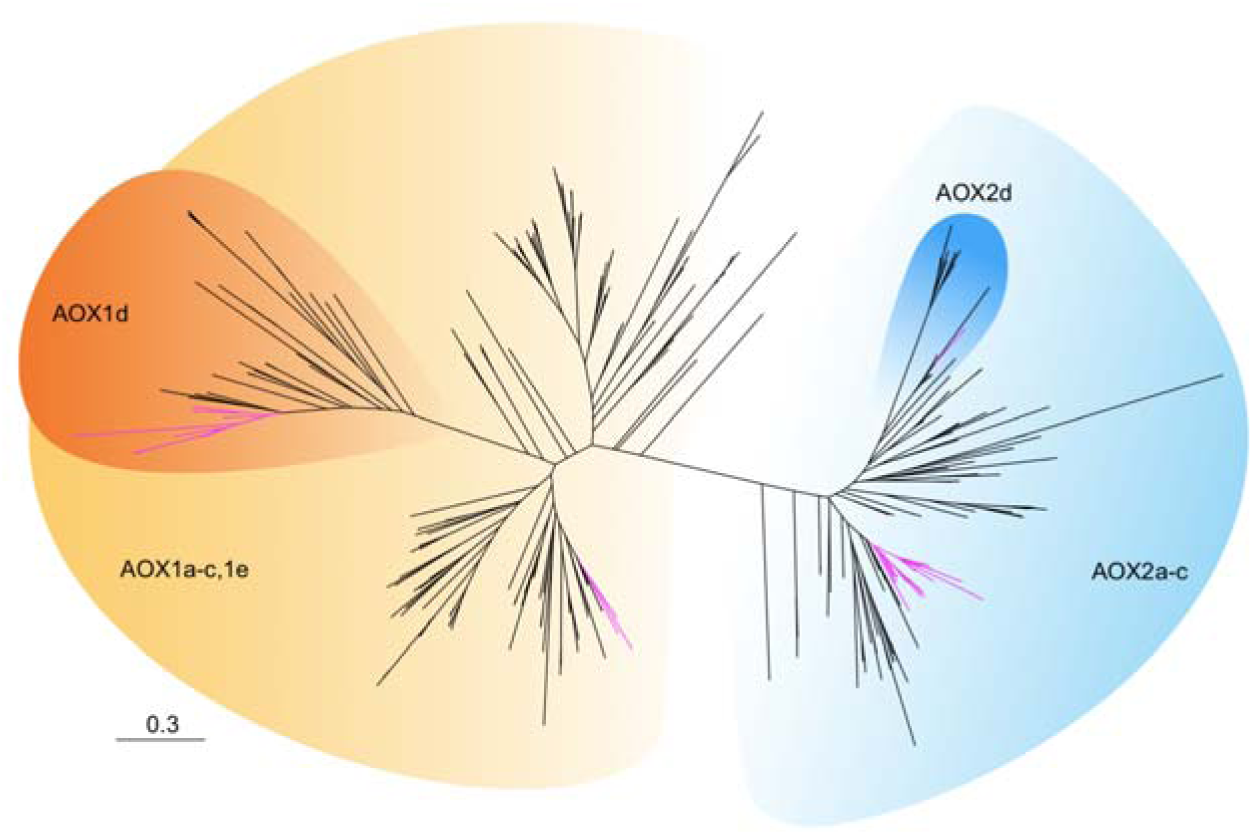
Phylogeny of AOX showing four major clades in angiosperms including AOX1a– c/1e, AOX1d, AOX2a–c, and AOX2d. The phylogeny was inferred using the Maximum Likelihood method implemented in IQTREE. Branches are drawn in proportion to genetic distance (substitutions per site) and Orobanchaceae gene copies are highlighted in magenta. Note the single Orobanchaceae branch in the AOX2d subclade resulting from a horizontal gene transfer from Rosaceae to Christisonia kwangtungensis. A fully annotated newick tree is provided in electronic supplementary material, data S1.

Increased reliance on alternative OXPHOS pathways in species with gene losses Despite our findings of significant N-mt OXPHOS gene loss, holoparasitic Orobancheae consistently demonstrated robust mitochondrial respiration, with similar OXPHOS efficiency compared to species showing no sign of gene loss (figure 3D; LME p-value = 0.612; electronic supplementary material, data S2). The normalized maximum respiration also showed a similar trend, with holoparasites having similar maximum respiration (figure 3B; LME p-value = 0.2146). Across individual complexes, holoparasitic Orobancheae tended to have higher FCFs in the fully nuclear-encoded complexes CII, AOX, and DH_ex_. However, this trend was only significant for CII, where holoparasitic Orobancheae demonstrated more than threefold higher CII flux (LME p-value = 0.02). Differences in the two alternative complexes DH_ex_ and AOX were not statistically significant (LME p-value >0.23), but showed a consistent trend of higher flux control in holoparasites by 21.3% and 39.1%, respectively (figure 3F, H). For the chimeric complexes CI and CIV, holoparasites had similar flux as the other taxa without gene loss (LME p-value >0.25).

**Figure 3.**
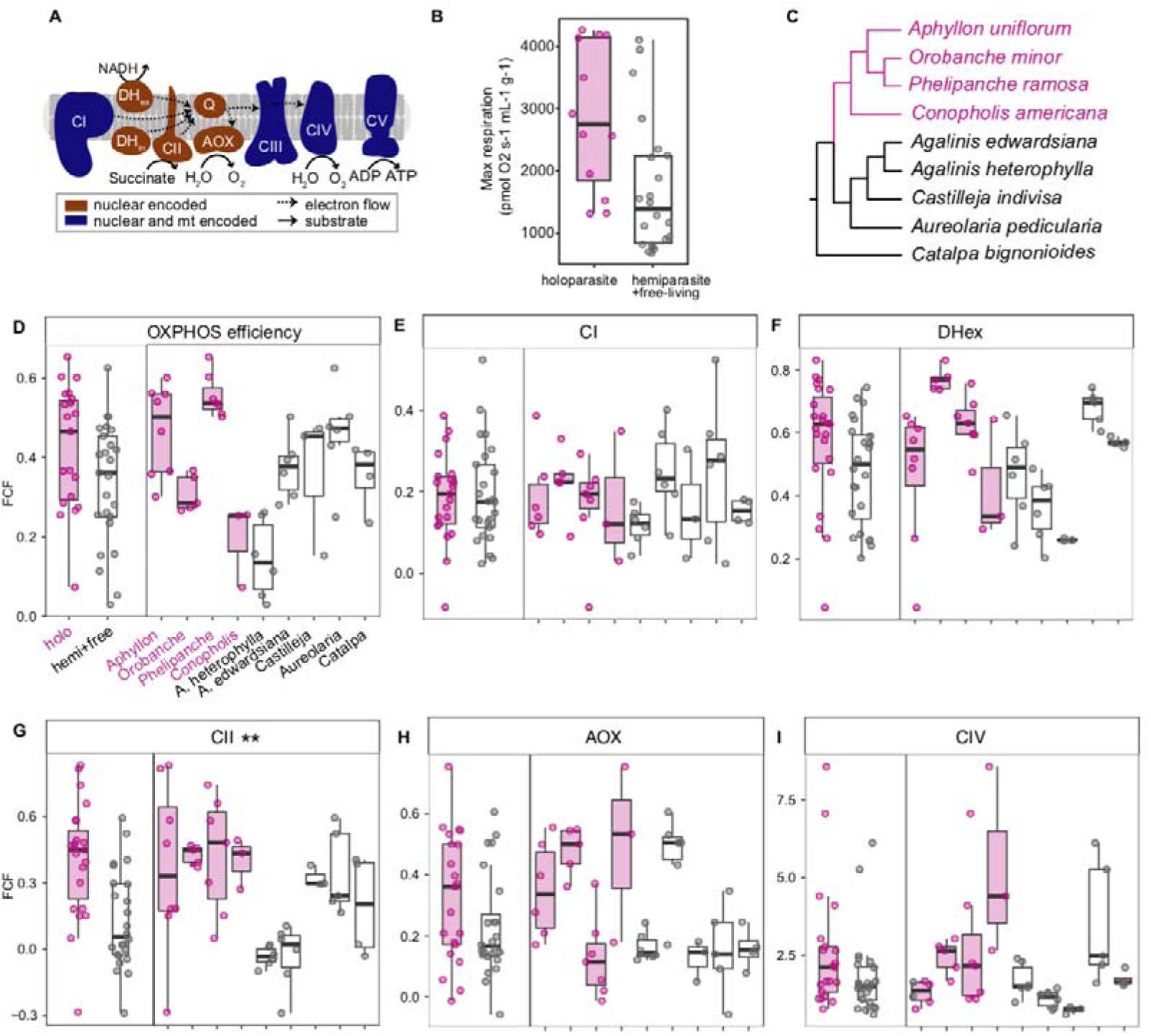
Increased reliance on nuclear-encoded OXPHOS pathways in holoparasitic Orobanchaceae. **A.** Plant mitochondrial electron transport chain components. Electrons enter through complex I (CI), CII, or the alternative NADH dehydrogenases (DH_ex/in_) and follow either the alternative oxidase pathway (AOX) or the cytochrome pathway CIII–CIV to reduce oxygen to water. CI, CIII, and CIV translocate protons to establish the proton motive force used by CV for ATP production. **B.** Normalized maximum mitochondrial respiration of holoparasitic species with extensive gene loss and their close relatives. **C.** Phylogenetic relationship among the nine species sampled in this study summarized from Mortimer et al. (Mortimer *et al*., 2022). **D-H.** Comparison of flux control factors (FCF) between the two groups of species for OXPHOS efficiency (**D**), CI (**E**), DH_ex_ (**H**), CII (**G**), AOX (**H**), and CIV (**I**). Statistical differences at p-value <0.05 are marked by ‘**’.

Overall, holoparasitic Orobancheae with gene loss tended to show higher flux levels across all measured variables, although only reaching statistical significance for CII (figure 3). However, this trend obscures considerable variation within and among species, and the overall trends observed between holo-versus hemiparasites are not uniformly distributed. For example, there was uniformly low AOX flux control in the holoparasitic Phelipanche ramosa and high AOX flux control in the hemiparasitic Agalinis edwardsiana (figure 3H), which goes against the general trend of high AOX flux in holoparasitic Orobancheae. Unfortunately, many of the species available for O2K experiments do not have available genetic resources, making it difficult to correlate putative losses of individual N-mt genes with OXPHOS flux.

Most FCFs did not show a significant correlation with each other, including comparisons among nuclear versus chimeric complexes (figure 4). However, a significant positive correlation was identified between the fully nuclear encoded DH_ex_ and CII FCFs (β = 0.241, LME *p*-value = 0.0057). Here, holo- and hemiparasites showed similar responses with the regression coefficient β being 0.242 and 0.207, respectively (figure 4A). None of the other pairwise comparisons, including nuclear versus chimeric (e.g., CI versus CII), chimeric versus chimeric (e.g., DH_ex_ versus AOX), or maximum respiration versus OXPHOS complexes (e.g., maximum respiration versus AOX), demonstrated significant global correlations (figure 4; electronic supplementary material, table S6). Some of the lack of correlation was likely driven by the disparate patterns shown in holo-versus hemiparasites. This was most obvious between maximum respiration rate and DH_ex_/CII flux (figure 4E-F). In both cases, the maximum respiration rates were negatively correlated with DH_ex_ (β = −2.2e3; LME p-value = 0.55) and CII (β = −4.4e3; LME *p*-value = 0.0083) in holoparasitic Orobancheae, but a positive correlation was recovered for DH_ex_ (β = 2.2e3; LME p-value = 0.08) and CII (β = 7.6e2; LME p-value = 0.37) in the hemiparasitic and free-living species.

**Figure 4.**
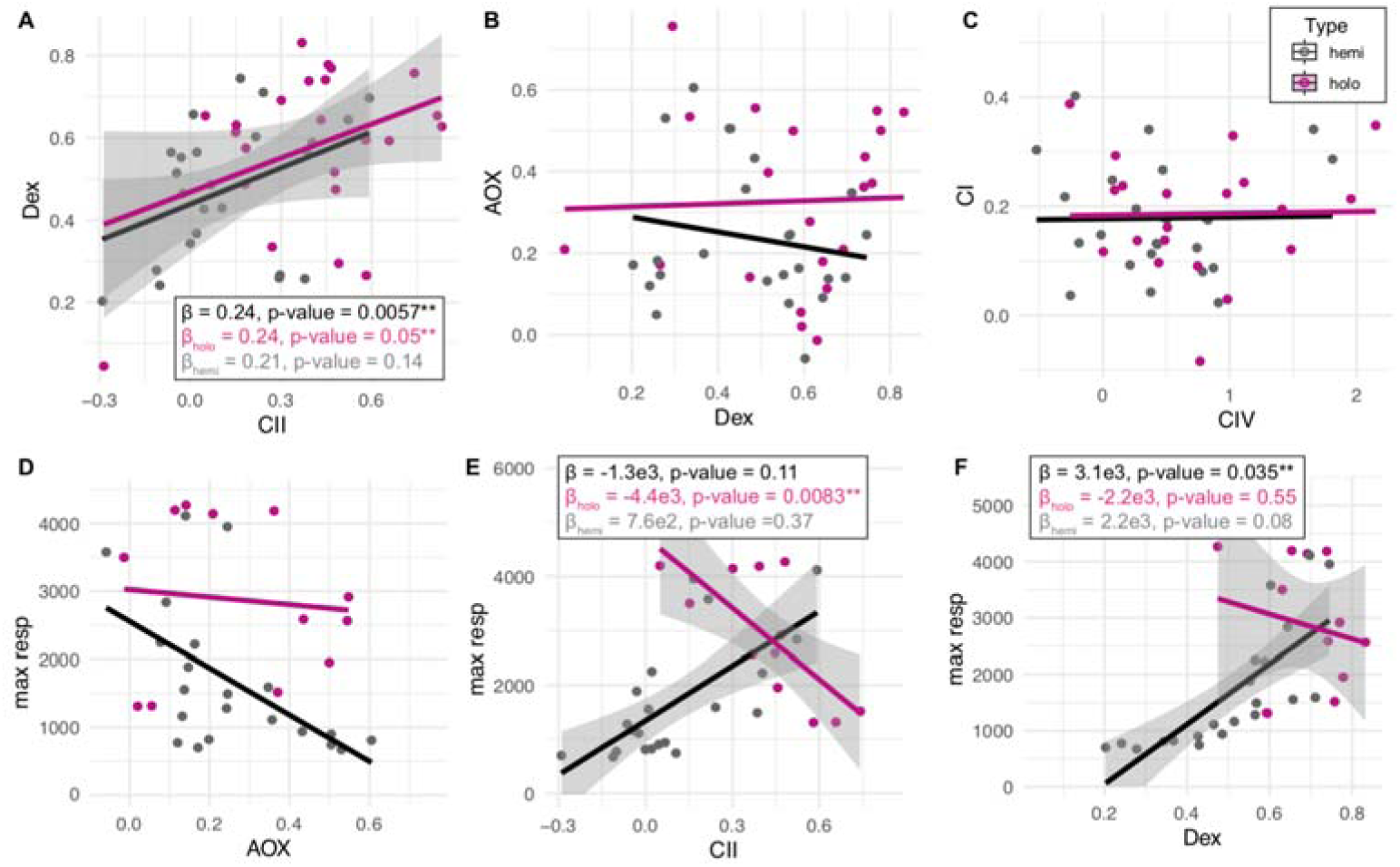
Relationships among FCF types in Orobanchaceae with different life history strategies. Magenta and black lines show the fitted linear mixed-effects (LME) models for holoparasitic and hemiparasitic Orobanchaceae, respectively. The standard deviation, p-values, and correlation coefficients β of LME models are only shown in significant relationships for the global (top) and partitioned datasets (middle and bottom).

## DISCUSSION

The parasitic lifestyle has been frequently associated with genome degradation in eukaryotes (Poulin and Randhawa, 2015). In land plants, it is clearly established that both the nuclear and plastid genomes are subject to gene loss and relaxed selection when parasitism evolves (Lyko and Wicke, 2021). Yet how parasitism or heterotrophism in general impacts the mitochondrial genome and function remains controversial. On one hand, unprecedented loss of the entire OXPHOS CI is documented in mistletoe and approximately 80% of the mitochondrial genes have been replaced by host-derived copies in some Balanophoraceae species (Skippington et al., 2015; Sanchez-Puerta et al., 2017). These massive losses and transfers are likely genetic hallmarks of relaxed selection due to cytonuclear conflict. On the other hand, mitochondria perform multiple critical housekeeping functions including ATP production and oxidative stress response that seem to be indispensable even for parasites. In both mistletoes and Balanophoraceae, the mitochondrial function under genetic impact is maintained by redundant or alternative pathways that are entirely nuclear-encoded (Gatica-Soria et al., 2024). In mistletoe, the production of ATP via glycolytic oxidation does not require mitochondria (Maclean *et al*., 2018). Our combined genetic and physiological investigation of Orobanchaceae mitochondria adds to the paradox that despite clear signatures of relaxed selection, respiratory functions can remain intact because of functional redundancy.

Unlike all other reported cases of mitochondrial gene loss in parasites, losses and transfers in Orobanchaceae are apparently restricted to nuclear-encoded genes. In our companion paper focusing on mitogenome evolution, all 45 species in Orobanchaceae displayed a complete and native set of core mitochondrial-encoded OXPHOS genes, including holoparasitic tribe Orobancheae (Cai *et al*., unpublished). The putative losses in N-mt genes are restricted to accessory subunits of OXPHOS complexes and account for no more than 20% of the total protein composition of each complex. Yet some of them may confer profound functional impact. For example, loss of the shared organellar RRR genes may lead to improper RRR machinery that promotes accelerated nucleotide substitutions (Parkinson *et al*., 2005; Guisinger *et al*., 2008; Sloan *et al*., 2012; Park *et al*., 2017). This was demonstrated in the plastid genome of *Geranium* (e.g., Parkinson *et al*., 2005) and is potentially relevant for the degenerated plastid genomes of holoparasites (Wicke *et al*., 2016). Widespread loss of the mitochondrial-specific RRR like *RECA3* in tribe Orobancheae may be linked with significantly accelerated mitochondrial genome shuffling we have found in these holoparasites (Cai *et al*., unpublished) because disruption of this gene resulted in extensive rearrangement of the mitochondrial genome in *Arabidopsis* (Miller-Messmer *et al*., 2012). Moreover, we found two putative losses of *ATP5* in *Aphyllon* and *Conopholis.* In *Arabidopsis,* reduced *ATP5* expression results in stunted seedling growth in dark conditions and altered leaf shapes in light due to lower ATP levels (Robison *et al*., 2009). In yeasts and *Fusarium* fungi, loss of *ATP5* similarly results in severely impaired nutritional growth (Uh *et al*., 1990; Yang *et al*., 2023). In Orobanchaceae, the parasitic lifestyle may allow species to cope with these otherwise deleterious mutations, but additional experiments and genomic data are needed to verify whether they are truly lost or expressed at low levels. For example, *ATP5* encodes the OSCP subunit of ATP synthase, which is sensitive to oligomycin. Therefore, lack of oligomycin sensitivity in *Aphyllon* and *Conopholis* would verify the unusual loss of *ATP5* in these species. Other losses can be confirmed by Western blots to examine the size of these protein complexes and explore the possibility of directly importing host-derived proteins or mRNAs. Demonstrating absence is challenging, but given the BUSCO evaluation, the biased phylogenetic distribution of missing genes, and the result from the RELAX analysis, we feel confident that relaxed selection on mitochondrial respiration has allowed the fixation of slightly deleterious mutations such as gene loss and horizontal gene transfer in the tribe Orobancheae.

At least eight putative horizontal gene transfers were detected in nuclear-encoded OXPHOS genes within Orobanchaceae, especially in *Christisonia*. Two-thirds of these genes are transferred from Poaceae, which are frequently invoked as both donors and receivers of horizontal gene transfers in previous investigations (Dunning *et al*., 2019; Hibdige *et al*., 2021). If these transfers are representative of the rest of the nuclear genome, then the 8.2% (n = 5) gene transfers make *Christisonia* the species most prone to nuclear gene transfers among land plants, which is nearly six times that of *Sapria* (Rafflesiaceae) at 1.4% across its nuclear genome (Cai *et al*., 2021). However, these gene transfers may be enriched in OXPHOS genes rather than reflecting the genomic average. Such enrichment of gene transfers in certain metabolic pathways aligns with the ‘functional necessity’ hypothesis of gene transfer where alien copies are recruited to compensate for the loss of indispensable functions such as mitochondrial respiration (Cai, 2023). Finally, one important caveat is that our data was derived from RNA sequencing and may represent host RNAs rather than gene transfers (David-Schwartz *et al*., 2008). Future genomic sequencing of holoparasitic Orobanchaceae including *Christisonia* will confirm the genetic background of these putative alien N-mt genes.

Despite widespread gene loss, horizontal gene transfers, and elevated non-synonymous substitutions indicative of relaxed selection, mitochondria of the holoparasitic Orobancheae performed similarly to the other species in high-resolution respirometry experiments, but tended to show a higher capacity for OXPHOS flux and overall mitochondrial respiration. However, intra-species variation was large even though these flux measurements were internally normalized. These variations could be caused by additional developmental and environmental factors that mobilized mitochondrial respiration in different ways. After accounting for these variations, the only significant difference we found was in CII capacity. Such a shift to increased CII flux may reflect a functional rescue of the compromised chimeric complexes due to gene loss, cytonuclear conflict, or overall dysfunction due to relaxed selection. For example, increased AOX respiration allows for adequate ATP production and maintenance of redox homeostasis in stressful environments (Vanlerberghe, 2013; Sweetman *et al*., 2019), and our study is in line with previous work where similar stress may be induced via molecular changes in mitochondrial genes (Weaver *et al*., 2020). Species in the holoparasitic tribe Orobancheae such as *Aphyllon* and *Orobanche* rely heavily on AOX and unlike other angiosperms, they may be using the more development-related AOX2 subclade for this purpose (Arnholdt-Schmitt *et al*., 2006; Costa *et al*., 2014). Even in hemiparasitic Orobanchaceae, the stress-related AOX1a-c/1e cannot be detected in most lineages and is replaced by the stress-related AOX1d subclade (Clifton *et al*., 2006). Future genomic investigation of Orobanchaceae will be able to confirm the differentiated retention of AOX subfamilies and their role in mitochondrial respiration.

We also found a positive correlation between CII and DH_ex_ (figure 4A), both of which are fully nuclear-encoded alternative electron entries to CI. The positive correlation between CII and DH_ex_ has been shown previously in *Silene* (Havird *et al*., 2019*b*; Weaver *et al*., 2020) and may suggest simultaneous mobilization of alternative complexes when CI is compromised. The disparate patterns between CII/DH_ex_ and maximum respiration in holo-versus hemiparasites (figure 4E-F) further corroborate the linked activities of alternative complexes and allude to the potentially different use of OXPHOS complexes in species with diverse life history strategies. Indeed, divergent signatures of selection are observed in both nuclear (electronic supplementary material, table S5) and mitochondrial-encoded genes (Cai *et al*., unpublished), with relaxed selection unique to holoparasitic Orobancheae.

Our results add to the growing consensus that in plants, compromised mitochondrial respiration originating from cytonuclear conflict may be compensated by elevated activities of the alternative pathways. However, there are drastically different mechanisms underlying cytonuclear conflict, leaving divergent genetic signatures in *Silene*, *Viscum*, and in our case, Orobanchaceae. For example, the exceptionally high substitution rate of mitochondrial genes in *Silene* is likely a result of cellular-level drift, especially in species with only one to two mitochondria per cell and possibly inefficient homologous repair mechanisms (Zwonitzer *et al*., 2024). These incompatible substitutions are at least partially compensated by both corresponding mutations in nuclear genes and physiological plasticity to alternative respiratory pathways (Sloan *et al*., 2014; Havird *et al*., 2017; Weaver *et al*., 2020). This mechanism is in direct contrast with the pattern observed in parasitic lineages, which is likely a result of relaxed selection on OXPHOS respiration. In the mistletoe *Viscum*, both nuclear and mitochondrial genes encoding CI have been lost and this functional deficiency is partially rescued by increased activities of AOX and alternative ATP generation through non-mitochondrial processes (Maclean *et al*., 2018; Senkler *et al*., 2018). In the holoparasitic Orobancheae, we found a general trend of relaxed selection for N-mt genes as well as their mitochondrial counterparts, but gene loss and horizontal gene transfer are restricted to N-mt genes (Cai *et al*., unpublished). Such biased rates of loss or transfer in the nucleus versus mitochondrion are also seen in the parasitic Balanophoraceae (Gatica-Soria *et al*., 2024) and potentially in Rafflesiaceae (Cai personal communication). This phenomenon may represent a lag time and evolution-in-progress where mutational changes are reflected in the faster-evolving nuclear genome before they are manifested in the mitochondrial genome, which is generally more slow-evolving in plants. Our research underscores the importance of studying parasitic plants as model systems to unravel fundamental questions about mitochondrial evolution and physiological adaptation to different life-history strategies.

## Supplementary Information Supplementary figures

### Supplementary figures

**Figure S1.**
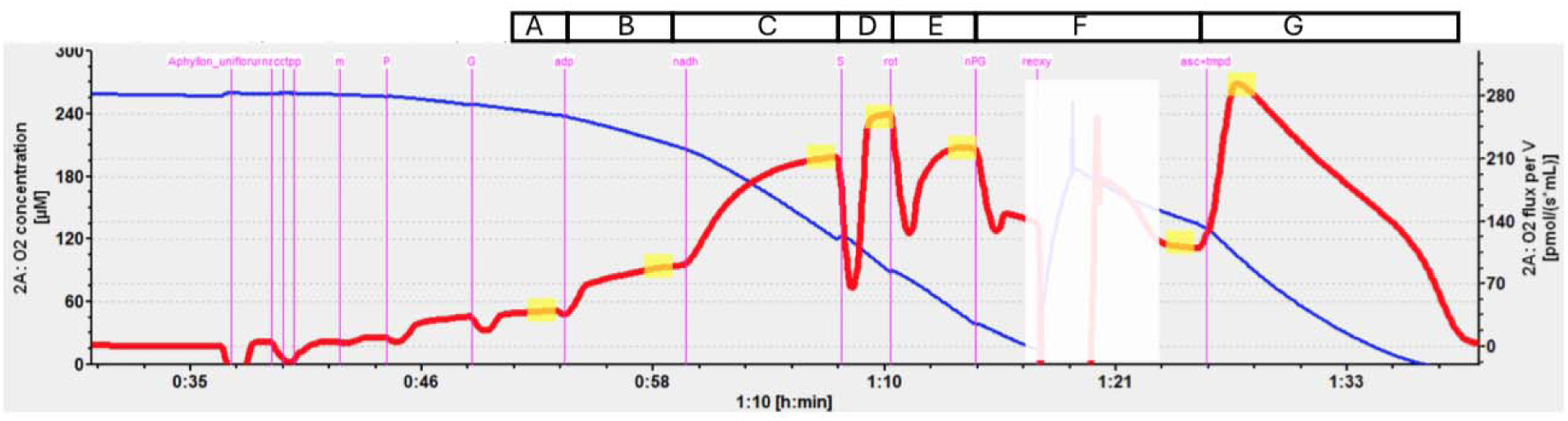
Quantification of mitochondrial respiration using O2K high-resolution respirometry. Data were taken from an Aphyllon uniflorum sample that showed typical changes in oxygen concentration (blue line) and oxygen flux (red line) by adding mitochondrial respiratory substrates or inhibitors over a 1.5-hour experiment period. Values of oxygen flux were recorded (highlighted yellow regions) after the addition of substrates or inhibitors (purple labeled text) for steps A-G as defined in Electronic supplementary material, Table S3. Reoxygenation events (in white box) were masked to enhance clarity.

### Supplementary tables

**Table S1** Taxon sampling, data source, and data quality evaluation for the genomic survey of nuclear-encoded mitochondrial genes in angiosperms.

**Table S2** Taxon sampling and collection localities for O2K high-resolution respirometry in Orobanchaceae.

**Table S3** Definition of respiratory flux control factors.

**Table S4** Pairwise sequence divergence to Arabidopsis thaliana inferred from concatenated sequence of nuclear-encoded mitochondrial genes.

**Table S5** Statistical significance of relaxed or intensified selection of mitochondrial protein complexes in Orobanchaceae.

**Table S6** Pairwise correlation between selected OXPHOS FCFs under the linear mixed-effects models.

Supplementary Notes

**Note S1** Plant mitochondrial isolation and O2K protocol

### Supplementary data

**Data S1** DNA codon alignments and maximum likelihood phylogeny for nuclear-encoded mitochondrial genes in angiosperm.

**Data S2** Raw oxygen flux and calculated flux control factors from fifty runs of O2K high-resolution respirometry in Orobanchaceae.

